# Uterine immune dysregulation after asynchronous transfer induced embryonic apoptosis in mice

**DOI:** 10.1101/2025.09.24.677934

**Authors:** Nana Yang, Hang Zhao, Shuyuan Sun, Jing Zhang, Meng Li, Xiangyun Li, Xinglong Wu

## Abstract

Asynchronous embryo transfer is widely used in assisted reproduction, yet its adverse effects and mechanisms remain unclear. This study investigates implantation failure in D1.5 pseudopregnant mice to elucidate mechanisms underlying asynchronous transfer. Blastocysts were transferred into uterine horns of D1.5 (Group A), D2.5 (Group B), and D3.5 (Group C) recipients. Implantation sites were counted at D7.5, and embryos were recovered within 3 hours post-transfer for analysis. Multibarcode single-cell RNA sequencing was applied to embryos and uterine tissues. Embryo recovery rate in Group A was significantly lower than in Groups B and C (*p*<0.01), with an abnormal embryo rate of 68.67%. Hoechst 33342/PI double staining revealed dead-cell features in abnormal embryos. Transcriptomic analysis of embryos showed downregulated DNA damage repair and structural maintenance genes in Group A. GO and KEGG analyses highlighted enrichment in transcriptional regulation and cell cycle pathways. GSEA identified activated apoptosis pathways, with intrinsic (mitochondrial) and extrinsic (death receptor TNFR-mediated) apoptosis genes significantly upregulated (*p*<0.05). Uterine transcriptome analysis revealed downregulated collagen family genes (ECM maintenance) but upregulated immune-related genes (IL family, CDs, C3) in Group A. Functional enrichment confirmed ECM and immune regulation pathway involvement. Gene network analysis identified *Clca1*, *Smpdl3a*, *Tmprss4*, *Muc4*, *Cfb,* and *Nupr1* as key regulators of D1.5 uterine immunity/inflammation. D1.5 uterine immune dysregulation induces embryonic stress and inflammation. Impaired DNA damage repair activates intrinsic apoptosis, while upregulated TNFR signaling triggers extrinsic apoptosis, collectively blocking embryonic development toward implantation competence. These findings provide mechanistic insights into optimizing embryo transfer timing and improving reproductive outcomes.

## INTRODUCTION

Embryo transfer technology has become an indispensable component in assisted reproductive technology and animal husbandry, particularly exhibiting significant value in the fields of in vitro fertilization and animal cloning. Embryo transfer in mice was first successfully conducted by McLaren in 1958^[1]^. Shortly after, Tarkowski successfully carried out the oviduct transfer of mouse embryos, marking a new era in the development of embryo transfer technology ^[2]^.

In the traditional embryo transfer technique, it was commonly believed that the developmental stage of the transferred embryo had to be synchronized with the uterine environment of the recipients. However, as research progressed, it was discovered that early embryos, from the fertilized egg to the blastocyst stage, could successfully implant even when transferred to the oviduct on day 0.5 of pseudopregnancy (D0.5) in mice ^[3]^. Some studies have confirmed that morula or blastocysts without a zona pellucida can also successfully implant after being transferred to the oviduct or uterus ^[4,5]^. Currently, asynchronous embryo transfer is not only used in mouse embryo transfer but also has been widely used in many different species, and has even become a common transfer method, including sheep ^[6]^, pigs ^[7,8]^, horses ^[9]^, single-humped camels ^[10]^, and humans ^[11]^.

Endometrial regulation of embryo development is a key factor influencing implantation in embryo transfer ^[12]^. A study found that when an asynchronous embryo was transferred into the reproductive tract of a recipient mouse, its development would be retarded when transferred to a ‘younger’ uterus. This pause continued until the implantation window in the recipient’s uterus opened ^[13,14]^. Embryonic development would be accelerated when embryos were transferred to ‘older’ uteri ^[15]^. Moreover, embryos at different development stages would implant at the same time ^[14,16]^. However, the study by Ueda et al. showed different results: the implantation of mouse embryos at different developmental stages, once transferred to the same recipient, was not synchronous. Furthermore, their early development after implantation also varied ^[17]^. Thus, regulating early embryo transfer into the uterus at different developmental stages is still controversial.

A comparison of pregnancy outcomes between asynchronous and synchronous embryo transfers shows that the demand for synchronicity donors and recipients varies considerably in different species. In rabbits, a 24-hour difference in asynchrony between donors and recipients did not affect implantation ^[18]^, but in humans, 24-hour asynchronicity decreased pregnancy rates after transfer from 20.5% to 11% ^[19]^. Noyes and Dickmann ^[20]^ reported that asynchronous transfer of rat blastocysts produced significantly heavier fetuses than synchronous transfer. Furthermore, mouse embryos survived considerably better when transferred to recipients that ovulated 24 hours after donors ^[21]^. Currently, transferring blastocysts or morulae to D2.5 and D3.5 uterus, or transferring various early-stage embryos to D0.5 oviduct are the commonly used methods for mouse embryo transfer ^[12]^. These results lead us to ponder whether the requirement for synchronicity of the physiological states between the donors and recipients in mouse transfer is not stringent. If a blastocyst was transferred into the D1.5 oviduct or uterus, can it be successfully implanted?

This study first attempted to transfer embryos to the D1.5 recipient uterus, but the embryos failed to implant successfully. Additionally, we were surprised to find a significant decrease in embryo quality when we recovered embryos from the uterus after the embryo transfer. This finding implies that transferred embryos may not develop to the window of implantation. Thus, this study transferred embryos to pseudopregnant mice on days 1.5, 2.5, and 3.5, then recovered embryos within 3 hours for transcriptome analysis. Additionally, transcriptome analysis was conducted on D1.5 to D3.5 recipient uterus tissue. Our findings aim to improve asynchronous transfer protocols and provide new insights and directions for studying the interaction between the uterine environment and embryonic development. Meanwhile, the study can also provide a certain theoretical basis for improving the pregnancy rate of human embryo transfer.

## MATERIALS AND METHODS

### Animals

SPF CD-1 mice (6–8 weeks old) were purchased from Beijing Vital River Laboratory Animal Technology Co., Ltd. (Beijing, China) and were housed in a stable environment (temperature, 22°C – 26°C; humidity, 50% – 60%; and a 12-h light/12-h dark cycle (i.e., lights on from 6 AM to 6 PM), with free access to food and water. All experimental protocols performed on the animals were approved by the Animal Care and Use Committee at Hebei Agricultural University (2021-011), China.

### Grouping and preparation

The presence of a vaginal plug after mating was designated as day 0.5 of pregnancy (Note: Typically, the prefix E is used for embryos, while the prefix D is used for days post-coitum or pseudopregnancy in females).

Preparation of donor mice: Female mice in natural estrus were selected and mated with male mice at a ratio of 1:1. On D3.5 post-mating, a disposable syringe equipped with a blunt-tipped needle was used to aspirate M2 medium and flush the blastocysts from the uterus of the donor mouse. The embryos were then cultured in M2 medium until they were transferred.

Preparation of recipient mice: To prepare the recipient mice, vasectomized male mice were paired with naturally cycling female mice at a ratio of 1:1. The female mice recipients were then divided into three groups: Group A-D1.5 recipients, Group B-D2.5 recipients, and Group C-D3.5 recipients. Embryo transfers were then performed on day 1.5, day 2.5, and day 3.5 after mating for each group.

### Embryo transfer

Before beginning the experiment, the recipient mice were anesthetized, and a small incision was made on the dorsal side of their bodies to expose the uterus. Eight to ten well-developed blastocysts were transferred into the right uterine horn of the D1.5, D2.5, and D3.5 recipient mice. The uterus was gently returned to its original position, and the incision was closed with sutures. The mice were then placed on a warming plate to recover from anesthesia. Afterward, the mice were individually placed in a new cage. The mice were euthanized by cervical dislocation and dissected to determine the number of embryo implantation sites on day 7.5. The embryo implantation rate equals the number of blastocysts transferred divided by the number of embryo implantation sites.

### Embryo collection and evaluation

After the embryos were transferred into the uterus of the recipient mice, the uterus was ligated on the side where the embryos were transferred to prevent embryo loss. Mice were euthanized by cervical dislocation at 1, 2, and 3 hours after embryo transfer, respectively. The uterus was removed and the junction between the oviduct and the uterus was cut. A sterile, disposable syringe was used to aspirate 0.2mL of M2 medium. A blunt-tipped 18-gauge needle was inserted into the cervix through the cervical os. The uterus was then flushed to obtain the embryos, and their status was observed. The embryo recovery rate is equal to the number of retrieved embryos divided by the number of transferred embryos. According to a simplified blastocyst grading system, the embryo quality is divided into three grades: Grade A - fully expanded with a clear inner cell mass and cohesive trophectoderm; Grade B- unexpanded with a clear inner cell mass and sticky trophectoderm; Grade C- small mass with irregular trophectoderm excluded or degenerate cells ^[22]^. The abnormal embryo rate is calculated by the number of abnormal embryos divided by the total number of recovered embryos.

### Sample preparation

Collect endometrial tissues from pseudopregnant mice on D1.5 (A; n=5), D2.5 (B; n=5), and D3.5 (C; n=5) respectively. Meanwhile, obtain embryos before transfer to the D1.5 uterus defined as pre-transfer embryos (D) and embryos before transfer to the D1.5 uterus defined as post-transfer embryos (E), ensure that each group (D and E) contains 2-3 embryos, with a total of 5 replicates per group (n=5). Prior to adding the embryos and endometrial tissues to the RNA lysis buffer labeled with a barcode, the zona pellucida must be removed using Tyrode’s solution (Sigma, Germany).

### RNA extraction, library construction and MultiBarcode single-cell transcriptome sequencing

Endometrial tissues and embryos with the zona pellucida removed (with a cell count of less than twenty thousand) are placed into lysis buffer containing barcode tags (RNase inhibitor, Triton X-100, dNTPs, barcode primer, nuclease-free water) for thorough lysis. All operations are carried out on ice. The lysis conditions are as follows: 25°C for 5 minutes, 42°C for 60 minutes, 50°C for 30 minutes, 72°C for 10 minutes, and then held at 4°C. Subsequently, reverse transcription is performed. The reverse transcription products are then added to a PCR amplification system consisting of KAPA Hifi Polymerase mix, IS PCR primer, 3’ end primer, and nuclease-free water for PCR amplification. The PCR program is set as follows: 95°C for 3 minutes, followed by 4 cycles of 98°C for 20 seconds, 65°C for 30 seconds, and 72°C for 5 minutes. Then, there are 10-16 cycles of 98°C for 20 seconds, 67°C for 15 seconds, and 72°C for 5 minutes, ending with a final cycle at 72°C for 5 minutes. The PCR products are purified using a Zymo kit and further purified twice with 0.8X AMPure XP beads. The final product is then dissolved in EB. The DNA fragments are then processed using the Zymo kit for DNA biotin enrichment. The enriched products are quality checked on a Qsep400, and the fragments are mainly distributed above 1 kb. The enriched cDNA fragments are then ligated to C1 beads using the KAPA Hyper Prep kit (KK8505) to construct a cDNA library. The library size is quality checked on a Qsep400, with fragments mainly distributed between 300 and 700 bp. Finally, Geekergene (Beijing, China) performs library sequencing on the Illumina Novaseq 6000 platform, generating 150 bp paired-end reads.

### Sequencing data quality control

After the sequencing process is completed, the raw sequencing data undergoes rigorous quality control (QC) to ensure the reliability and accuracy of the results. Through Perl scripts, we can filter the following types of unqualified data:1) TSO sequences; 2) polyA sequences; 3) Sequences with a high percentage of which are the bases that were not measured due to machine equipment or other reasons; 4) Sequences with junctions that have not been cleanly removed or have been punctured due to short read lengths; 5) Short fragments, which are sequences that are less than 37bp after removing the splice; 6) Low quality sequencing, which refers to sequences with low quality bases.

### Sample gene quantification

Expression matrices were generated by aligning reads with the mm10 reference genome using STAR (v2.7.9a). The number of genes per cell, mitochondrial content and other metrics were obtained with the help of Seurat. Samples with intragroup duplicates were then analyzed for differential expression using DEseq2. Genes with a false discovery rate (FDR) < 0.05 and an absolute fold change (FC)> 2 were considered differentially expressed genes (DEGs).

### Gene Ontology and Kyoto Encyclopedia of genes and genomes analyses

All DEGs were analyzed using clusterProfiler against the Gene Ontology (GO) database (http://www.geneontology.org/) to identify enrichment in three parts: cellular component (CC), molecular function (MF) and biological process (BP). The Kyoto Encyclopedia of Genes and Genomes (KEGG, http://www.genome.jp/kegg/) pathway enrichment analysis was compared with the whole genome (*p*<0.05).

### Gene set enrichment analysis

Gene Set Enrichment Analysis (GSEA) is a method that identifies whether a predefined set of genes (i.e., a gene set) exhibits statistically significant and consistent differences between two biological states (or two groups). The plots were drawn based on R version 4.1.3 (2022-03-10) on the OmicStudio platform (https://www.omicstudio.cn).

### Standardization of expression mean

STEM (Standardization of Expression Mean) analysis is a powerful method for identifying genes that exhibit similar patterns of change within a given dataset. We utilized the OmicShare online platform (https://www.omicshare.com/tools/) to perform groups A, B, and C analysis, which classify of genes based on their expression trends. *p*<0.05 was considered statistically significant, and the number of trends was chosen to be 20.

### Weighted gene co-expression network analysis

WGCNA (Weighted gene co-expression network analysis) is a common method to construct a gene co-expression network. Each node is defined as a gene, and genes with common expression in different samples are in the same gene network, and the expression correlation coefficient between them generally measures the co-expression relationship between genes. We used the SolarGenomics online platform (SolarGenomics.com) to perform WGCNA on genes with RPKM >1 and coefficient of variation (cv) of 0.5 or higher, which were obtained from transcriptome sequencing of uterine samples at different stages. Genes with kME > 0.7 are selected as module members (kME is a value used to assess the effective connectivity between key genes). Additionally, we employed Cytoscape software to identify key genes within the identified modules.

### Embryo staining

The double staining procedure using Hoechst 33342 (Invitrogen®, Shanghai, China) and Propidium Iodide (PI; Yeasen, Shanghai, China) is outlined as follows: Initially, the PI solution, which was originally at a concentration of 1mg/mL, was diluted to a ratio of 1:250.

The embryos were then subjected to the diluted PI solution for a staining period of 5 min in the dark to avoid photobleaching. Following this step, the embryos were thoroughly rinsed with PBS 3 to 5 times to remove excess stains. Subsequently, the embryos were stained with Hoechst 33342 at a concentration of 10μg/mL for 10 min. After completion, the embryos were once again washed with PBS for 3 to 5 times. Finally, the stained embryos were carefully mounted onto a microscope slide and observed under an inverted microscope (Nikon, Japan) for analysis.

### Data statistics and analysis

Except for sequencing data, all experimental data were analyzed for significance using an unpaired t-test with the help of GraphPad Prism 9.0. The results were expressed as mean±SEM. A *p*-value of less than 0.05 indicates a significant difference, while a *p*-value of less than 0.01 indicates an extremely significant difference. Sequencing data were expressed as mean FPKM value ±SEM and analyzed using the OmicStudio Cloud Platform (https://www.omicstudio.cn/tool).

## RESULTS

### Blastocyst transfer into D1.5 recipient uterus fails to implantation

About 8 well-developed blastocysts were transferred into recipient uteri on D1.5 (group A), D2.5 (group B), and D3.5 (group C) to compare implantation outcomes. It was observed that there was no significant difference in implantation rate between group B and group C (*p*>0.05). However, group A showed unsuccessful implantation after embryo transfer compared to groups B and C (*p*<0.01; Figure 1 and Table 1).

**Figure 1.**
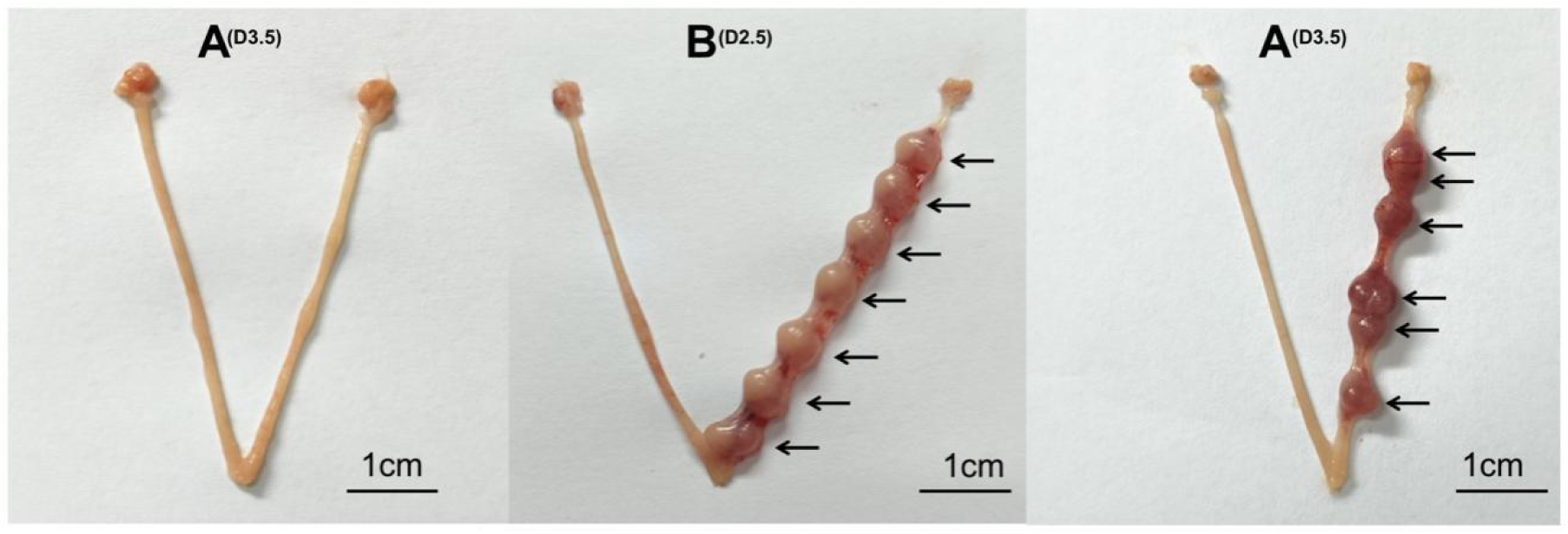
Implantation sites of embryos in the three groups were detected on Day7.5. Eight embryos were transferred into each recipient’s uterus. A, D1.5 recipients; B, D2.5 recipients; C, D3.5 recipients. Black arrows indicate the implantation sites (bar = 1 cm).

**Table 1.** The embryonic implantation of blastocysts transferred into different pseudopregnant recipients on Day7.5.

| *Groups* | *Total number of*<br>*transferred embryos*<br>*(n)* | *Number of*<br>*recipients(n)* | *Total numbers of*<br>*embryo implantation*<br>*sites(n)* | *Embryo*<br>*implantation rate*<br>*(%)* |
| --- | --- | --- | --- | --- |
| A | 63 | 8 | 0 | 0<sup>B</sup> |
| B | 64 | 8 | 45 | 70.31±1.86<sup>A</sup> |
| C | 61 | 8 | 40 | 65.85±0.58<sup>A</sup> |

*Note:* Different superscript uppercase letters in the same column designate a significant difference ( $p < 0.01$ ). The same capital letter means the difference is not significant ( $p > 0.05$ ). Values are the mean ± SEM based on 8 replicates per group. Implantation sites were counted in all groups on day 7.5. The embryo implantation rate is calculated as the number of blastocysts transferred divided by the number of embryo implantation sites. A, D1.5 recipients; B, D2.5 recipients; C, D3.5 recipients.

### Blastocyst of transfers into the D1.5 recipient uterus is gradually deteriorating

To determine the reasons for implantation failure, we transferred 8-10 blastocysts into the uteri of D1.5, D2.5, and D3.5 recipient mice and recovered embryos at 1h, 2h, and 3h after embryo transfer. As shown in Fig. 2A, unlike the embryos transferred into D2.5 and D3.5 recipient mice (which remained in good condition), the blastocysts transferred into D1.5 recipient mice gradually showed adverse changes over time. These blastocysts showed significant embryo retraction, a marked expansion of the perivitelline space, and even wholly lost the zona pellucida in some embryos (Fig. 2A). These morphological changes directly reflected a sharp decline in embryo quality. The statistical analysis showed that the number of embryos recovered from group A was significantly lower than those from groups B and C. The proportion of grade C embryos was significantly higher in group A than in groups B and C (*p*<0.01; Table 2). Furthermore, Hoechst 33342 and PI double staining showed that the cells in post-transfer blastocysts (group E) displayed prominent staining features characteristic of dead cells (Fig. 2B). These results indicated that cell death occurred in the embryos after they were transferred to the D1.5 uterus and could not develop to the window of implantation.

**Figure 2.**
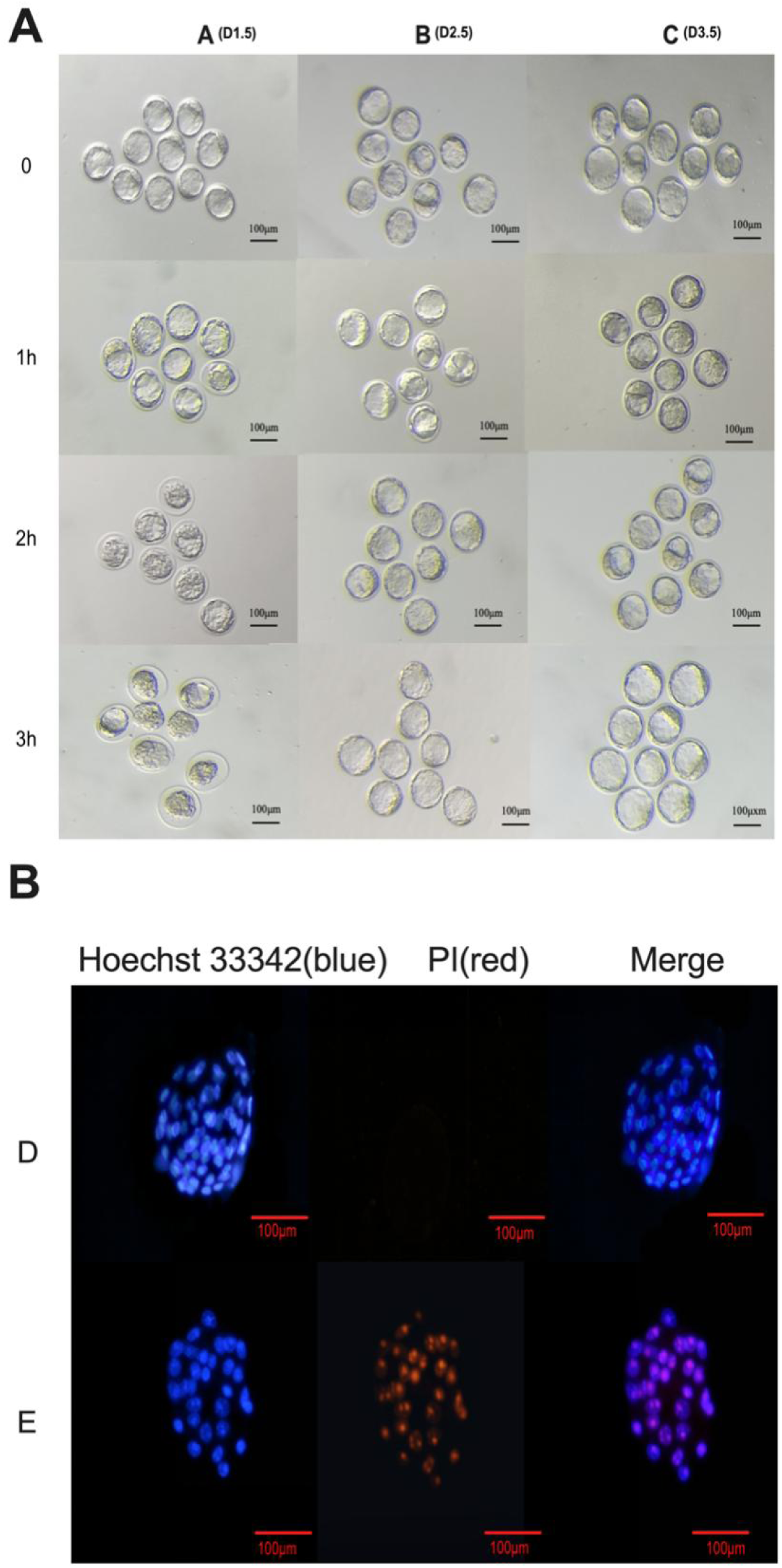
Development of embryos and embryo staining. (A) Development of embryos after 0 to 3h transfer to different recipient uteri. Ten embryos were transferred into each recipient’s uterus. A, D1.5 recipients; B, D2.5 recipients; C, D3.5 recipients. (bar = 100μm). (B) Hoechst 33342 and Propidium Iodide (PI) double immunofluorescence of embryos (200X magnification). Hoechst 33342 stains nuclei blue; PI stains dead cells or nuclei red. Scale bar: 100 μm. Note: Representative images are shown from Groups D and E.

**Table 2.** Recovery rate of blastocysts after 3h of transferring into different recipient.

| *Groups* | *Total number of transferred embryos (n)* | *Number of recipients(n)* | *Embryo recovery rate (%)* | *Abnormal embryo rate (%)* |
| --- | --- | --- | --- | --- |
| A | 125 | 15 | 56.02 ±5.85<sup>C</sup> | 68.67±6.30<sup>A</sup> |
| B | 120 | 15 | 74.85±4.07<sup>B</sup> | 11.49±3.07<sup>B</sup> |
| C | 124 | 15 | 89.81±2.09<sup>A</sup> | 7.98±2.06<sup>B</sup> |

*Note:* Different superscript uppercase letters in the same column designate a significant difference ( $p < 0.01$ ). The same capital letter means the difference is not significant ( $p > 0.05$ ). Values are the mean ± SEM based on 15 replicates per group. All embryos were recovered and assessed 3 hours after transfer into the recipient uteruses. The embryo recovery rate is calculated as the number of retrieved embryos divided by the number of transferred embryos. The abnormal embryo rate is calculated by dividing the number of abnormal embryos by the total number of recovered embryos A, D1.5 recipients; B, D2.5 recipients; C, D3.5 recipients.

### The apoptosis pathway of embryos transferred into the D1.5 uterus is activated

RNA sequencing was performed to investigate changes in embryo and uterus transcriptional profiles. The raw data from sequencing was filtered for quality control using Perl scripts. Both uterus and embryo samples had an error rate of less than 0.05% and a clean rate of over 85%, indicating excellent sequencing quality (Table 3). Principal component analysis (PCA) of transcriptomes showed clear separation groups D and E (Fig. 3A). The correlation heatmap displayed a significant positive correlation between individual embryo samples (Fig. 3B). Screening for DEGs with the criteria of FDR < 0.05 and |FC| > 2. When comparing group D with group E of embryos, 215 DEGs were significantly up-regulated, and 209 DEGs were significantly down-regulated (Fig. 3C). GO enrichment analysis showed the DEGs were mainly enriched in processes related to transcriptional regulation and the cell cycle (Fig. 3D). The bubble diagram showed that the top 20 enriched pathways included basal transcription factors, apoptosis, lysosomes, and TNF signaling pathways, etc (Fig. 3E). GSEA analysis indicated that the basal transcription factors pathway was inhibited in group E. Conversely, the apoptosis, lysosomes, and TNF signaling pathways were activated (Fig. 4A-D). Further analysis of the major DEGs associated with these pathways revealed that both *Atf4*, *Fos, Jun* and *Ctsb* were linked to pro-apoptotic processes. Additionally, the upregulation of *Gadd45g* and Atf4 was associated with stress responses, leading to the speculation that they could be involved in the activation of cellular apoptosis pathways. (Fig. 4F-H). In contrast, *TAFs*, which were involved in basal transcription and function as coactivators was significantly down-regulated (Fig. 4E). Compare the FPKM values of genes related to intrinsic and extrinsic apoptotic pathways to investigate specific apoptotic pathways further. The results indicated that the intrinsic pro-apoptotic factors *Bax* and *Bad* were significantly up-regulated in Group E (*p*<0.05). Similarly, extrinsic apoptosis-related genes *Casp-3*, *Casp-7*, and members of the TNF-receptor superfamily, including *Tnfrsf11a*, *Tnfrsf12a*, *Fas*, and CD40, were also highly expressed (*p*<0.05; Table 4). These findings suggested that apoptosis occurred in the embryos after their transfer to the D1.5 uterus.

**Figure 3.**
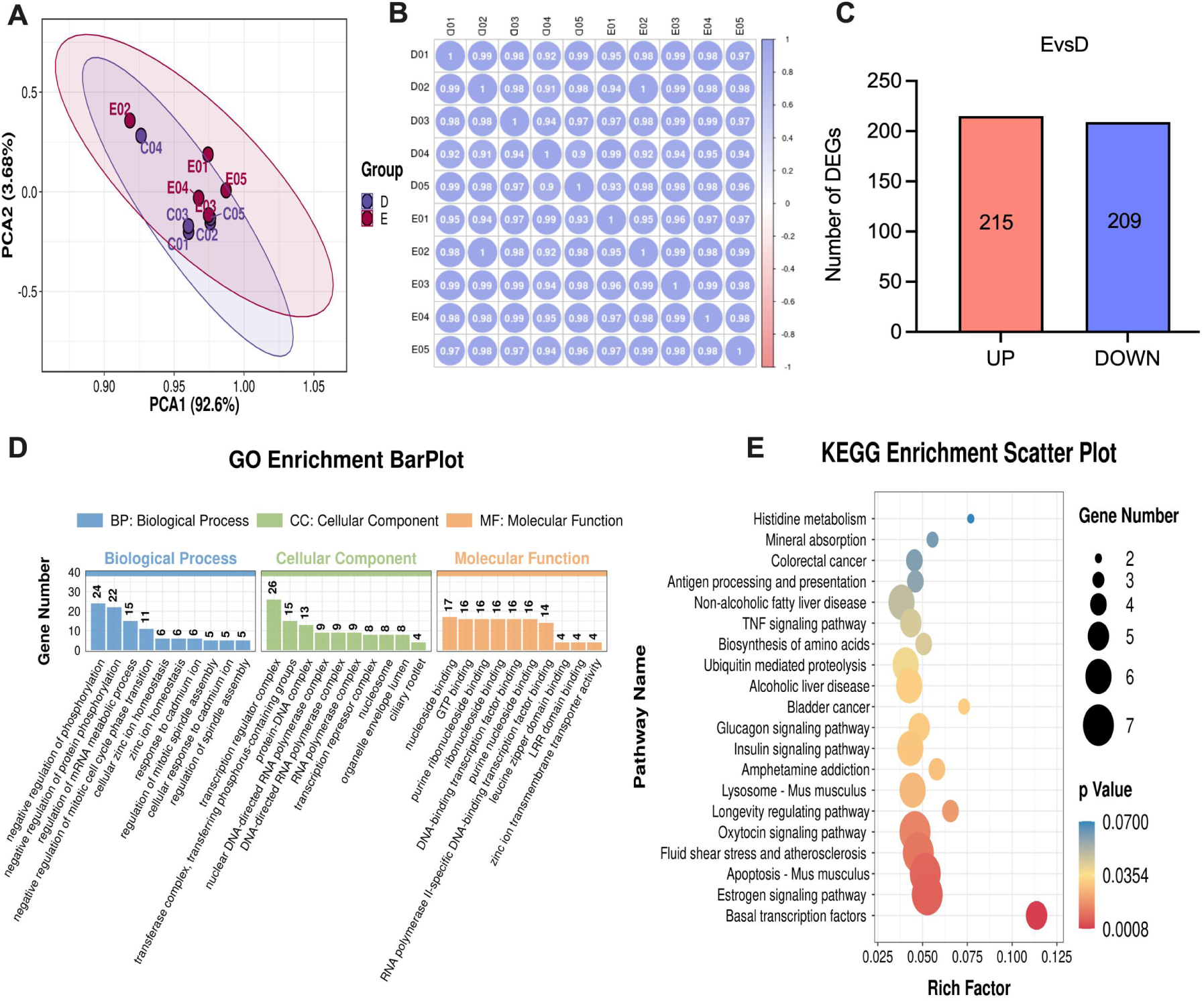
Transcriptome differences in embryos. (A) Principal component analysis (PCA) of embryos samples. (B) Correlation heat map between groups D and E. (C) Bar plots of differentially expressed genes (DEGs) in groups D and E. Red represents up-regulated DEGs; blue represents down-regulated DEGs. (D) The top 20 of Gene Ontology (GO) analysis of DEGs between groups D and E. Each colored bar represents a different process and the height of the bar indicates the number of DEGs. (E) The top 20 of Kyoto Encyclopedia of Genes and Genomes (KEGG) pathway analysis of DEGs between groups D and E. The color of the dot signifies the *p*-value of the hypergeometric test, where smaller values indicate greater reliability and statistical significance of the test. The size of the dot represents the number of genes; the larger the dot, the more DEGs in the pathway.D, pre-transfer blastocysts; E, post-transfer blastocysts. (n = 5).

**Figure 4.**
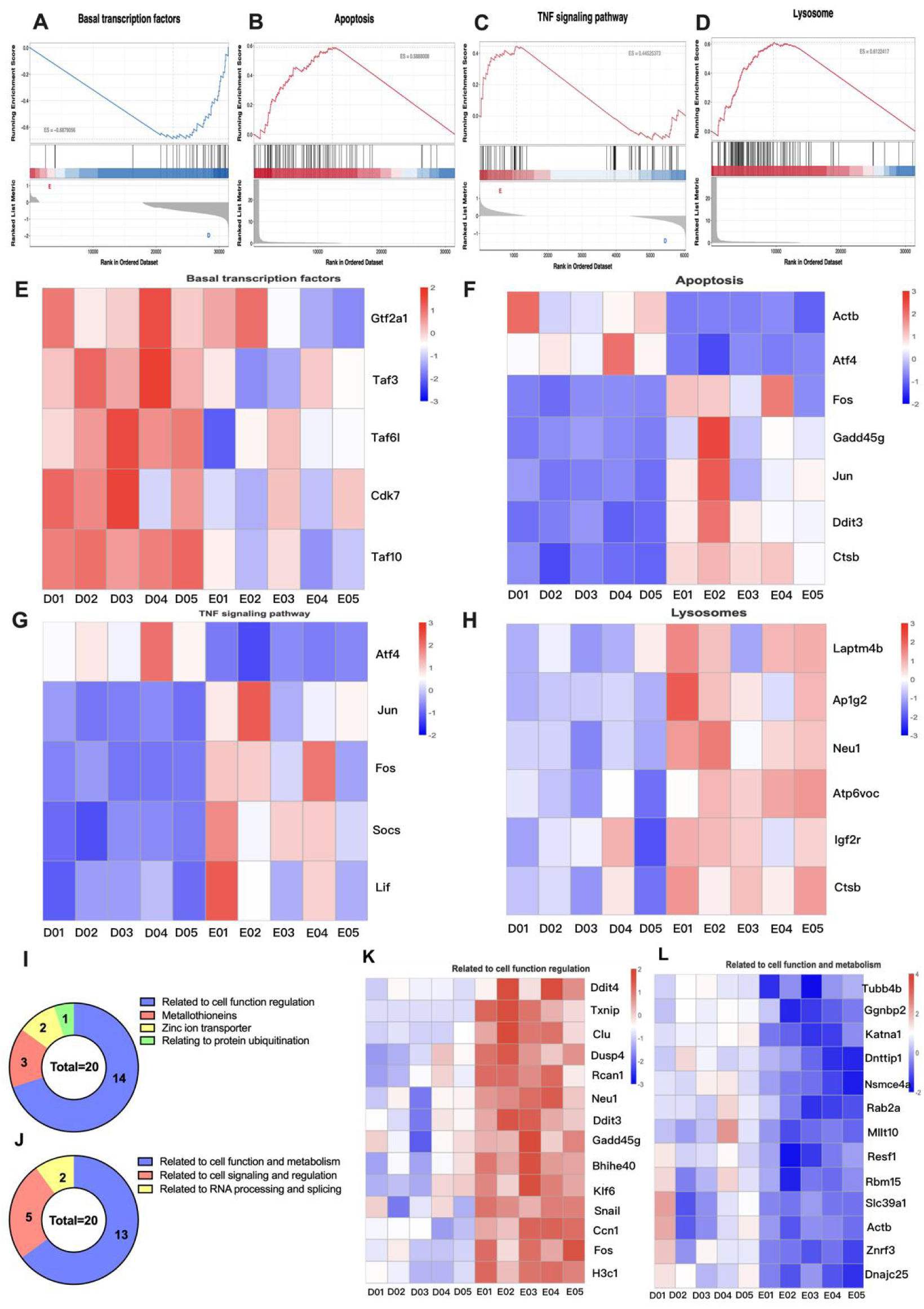
Abnormal transcription of post-transfer blastocysts. (A-D) Gene Set Enrichment Analysis (GSEA) maps and Heatmaps of basal transcription factors, apoptosis, TNF signaling pathway, lysosomes between groups D and E. The GSEA plot consists of three main parts: 1) The horizontal axis represents each gene in the expression dataset, and the vertical axis represents the corresponding enrichment score (ES). The peak value represents the maximum ES value of the gene set. When the running enrichment score (RES) plot is negative, the gene to the right of the peak is considered the core gene; when the RES plot is positive, the gene to the left of the peak is the core gene. The positive and negative ES values can be used to judge whether the pathway is activated or inhibited in the corresponding group. For example, in this figure C, compared with group D, ES = −0.687, indicating that basal transcription factors are inhibited in group E. 2) The second part is the gene location map. The expression dataset is arranged according to the expression abundance, and the location of the genes appearing in the functional gene dataset is indicated by the black vertical lines. 3) The third part is the distribution of rank values of all genes. (E) Heatmaps of basal transcription factors between groups D and E. (F) Heatmaps of apoptosis between groups D and E. (G) Heatmaps of TNF signaling pathway between groups D and E. (H) Heatmaps of lysosomes between groups D and E. For the heat maps, red represents up-regulated DEGs; blue represents down-regulated DEGs. (I) Donut chart of top 20 up-regulated and down-regulated (J) DEGs in group E. Different colors represent different gene function classifications in the Donut chart, and different numbers represent the number of DEGs in each functional category. (K) Heat maps related to cell function regulation and (L) related to cell function and metabolism between groups D and E. For the heat maps, red represents up-regulated DEGs; blue represents down-regulated DEGs. D, pre-transfer blastocysts; E, post-transfer blastocysts. (n=5).

**Table 3.** Quality control of embryos and uteri after sequencing.

*Note:*24-hbn-z001 & 24-hbn-z002; Naming of uteri samples for transcriptome quality
| *Sample* | *Raw reads* | *Raw_*<br>*bases* | *Clean*<br>*reads* | *Clean_base*<br>*s* | *Clean*<br>*Rate* | *Error*<br>*Rate* | *Q20* | *Q30* | *GC_*<br>*content* |
| --- | --- | --- | --- | --- | --- | --- | --- | --- | --- |
| *24-hbn-z001* | *38540103* | *5.781e*<br>*+09* | *38206987* | *5.060e+09* | *87.53* | *0.04* | *98.37* | *94.89* | *45.61* |
| *24-hbn-z002* | *63881940* | *9.582e*<br>*+09* | *63014195* | *8.589e+09* | *89.62* | *0.0414* | *96.81* | *91.64* | *46.15* |
| *hbn-z002-NT* | *29932584* | *44898*<br>*87600* | *29600991* | *3890334270* | *86.6271* | *0.04* | *98.055* | *94.53* | *45.395* |

control;
hbn-z002-NT; Naming of embryos samples for transcriptome quality control;
*Q20*; the percentage of bases with a Quality Score greater than or equal to 20 during sequencing. This Quality Score value is based on the Phred Quality Scoring System and measures the probability that each base is correctly identified. Specifically, Q20 corresponds to a 1% chance of misidentification, i.e., for every 100 bases, 1 base is expected to be misidentified.
*Q30*; percentage of bases with a quality value greater than or equal to 30 in the sequencing data.

**Table 4.**
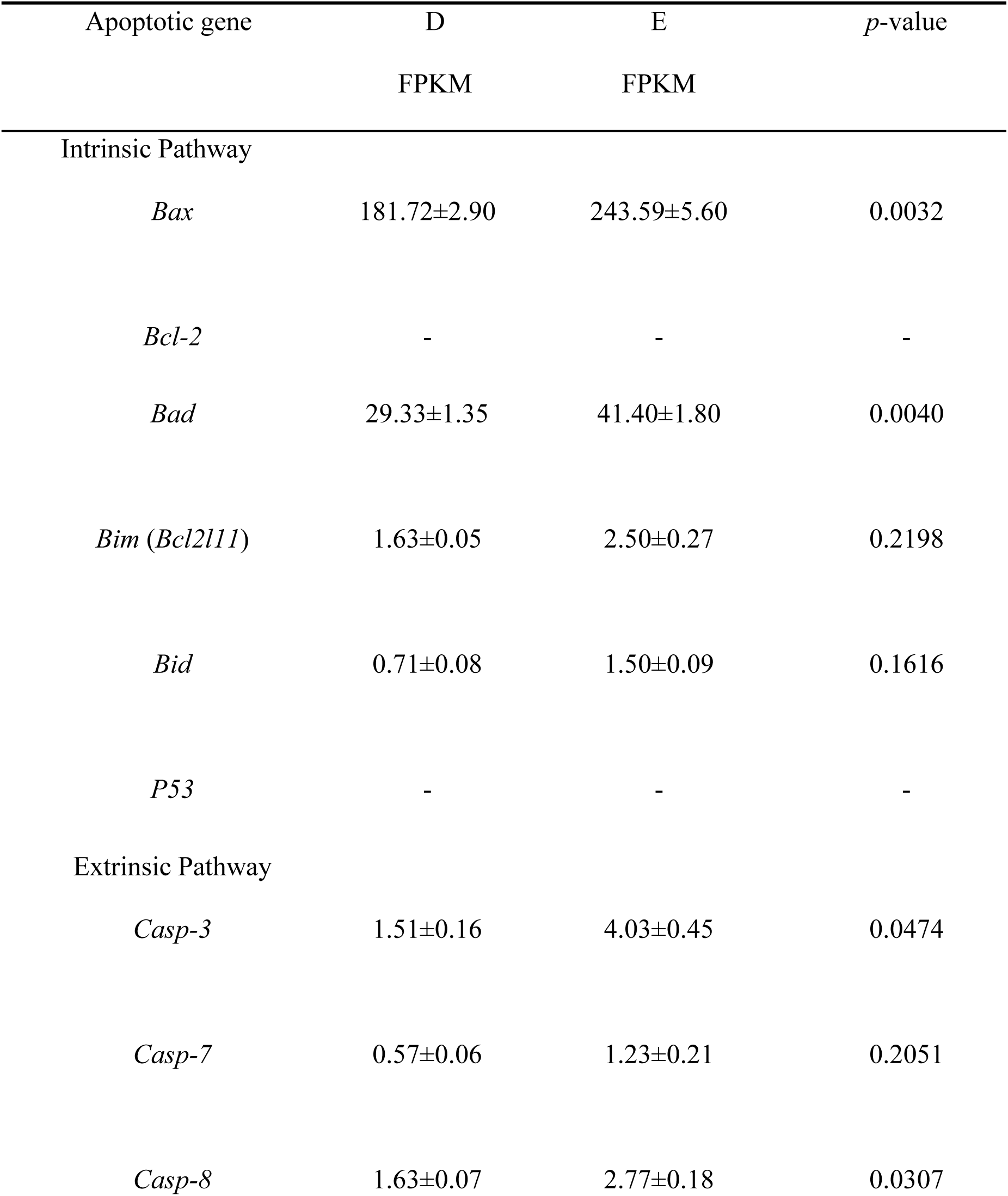
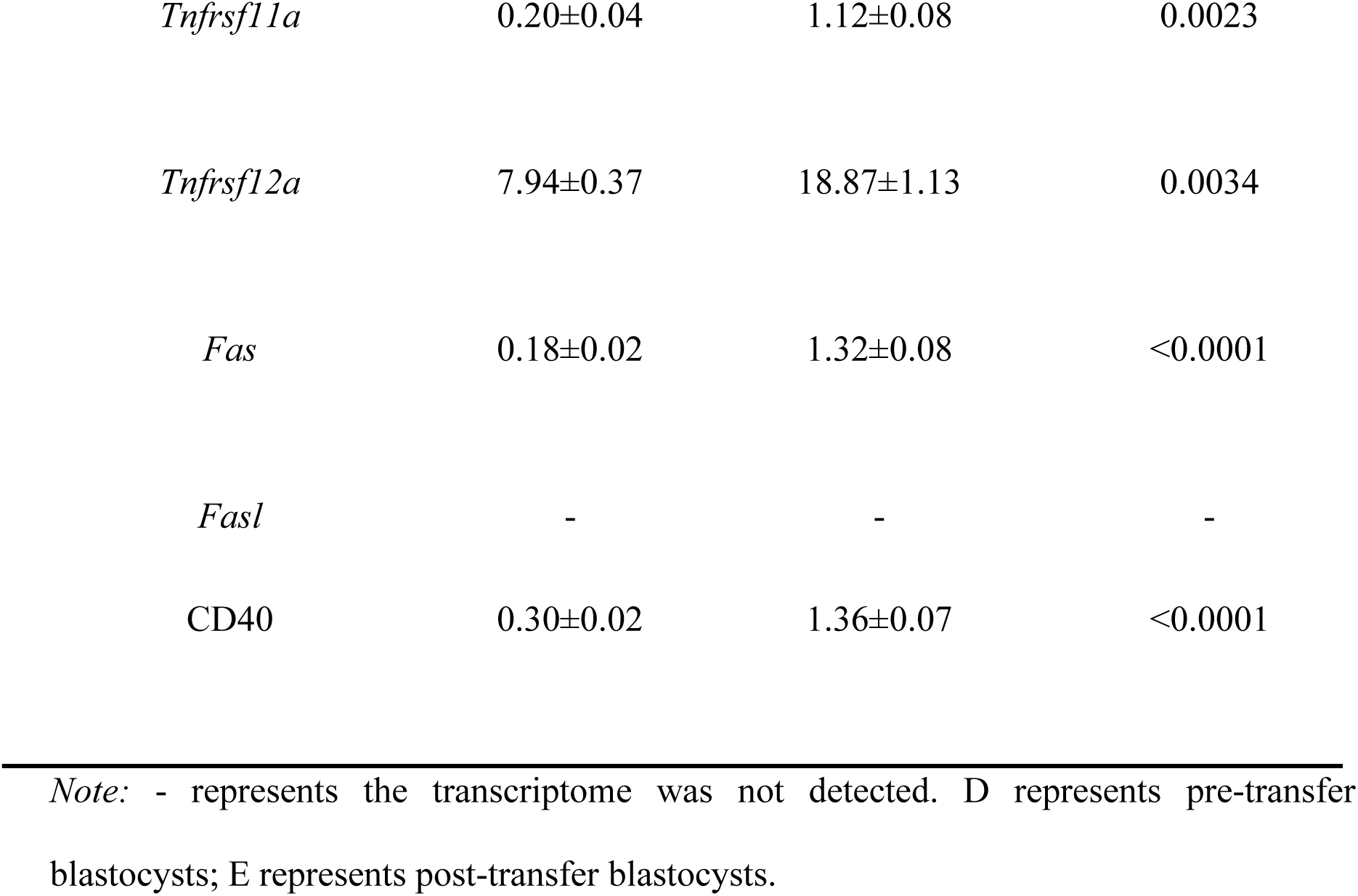
The FPKM level of apoptotic gene between groups D and E.

### DNA damage induces apoptosis in blastocysts once they are transferred to the D1.5 **uterus**

The intrinsic apoptotic pathway is primarily activated due to unrepairable DNA damage. The top 20 significantly up-regulated and down-regulated genes in post-transfer blastocysts were analyzed to verify whether the intrinsic pathway was activated. The top 20 up-regulated genes could be classified according to their functions as follows: metallothioneins (3/20), zinc ion transporter (2/20), related to cell function regulation (14/20), and related to protein ubiquitination (1/20) (Fig. 4I). The top 20 down-regulated genes were mainly related to RNA processing and splicing (2/20), cell signaling and regulation (5/20), and cell function and metabolism (13/20) (Fig. 4J). The heat map showed that among the up-regulated genes related to cell function and regulation, *Clu, Ddit4, Gadd45g*, and *Fos* were related to the stress response in group E (Fig. 4K). Among the genes related to cell function and metabolism, *Nsmce4a* was related to the repair of cellular DNA damage, and *Znrf3* was related to the maintenance of cellular structure. Meanwhile, the levels of *Znrf3, Slc39a1, Rab2a, Actb*, and *Tubb4b* were significantly down-regulated in group E (Fig. 4L). All of these DEGs were related to cell structure maintenance and DNA damage. Thus, these results suggest that embryo apoptosis is caused by DNA damage after their transfer to the D1.5 uterus.

### The structural stability of D1.5 uterine tissue decreases

The uterus is essential for embryo development, so transcriptomic changes in the uterus were analyzed by RNA sequencing. Principal component analysis (PCA) of transcriptomes showed clear separation among all groups (Fig. 5A). The correlation heatmap displayed a significant positive correlation between individual samples, with the D1.5 uterus sample showing significant differences from the other samples (Fig. 5B). A total of 2577 DEGs were identified comparing group A with group B, with 1179 DEGs up-regulated and 1398 DEGs down-regulated. The comparison between groups A and C yielded 2615 DEGs, including 1379 up-regulated and 1236 down-regulated genes (Fig. 5C). Furthermore, 958 DEGs were found to be exclusively up-regulated in group A, while 892 DEGs were exclusively down-regulated in group A (Fig. 5D). These data indicated significant differences between uteri at different stages.

**Figure 5.**
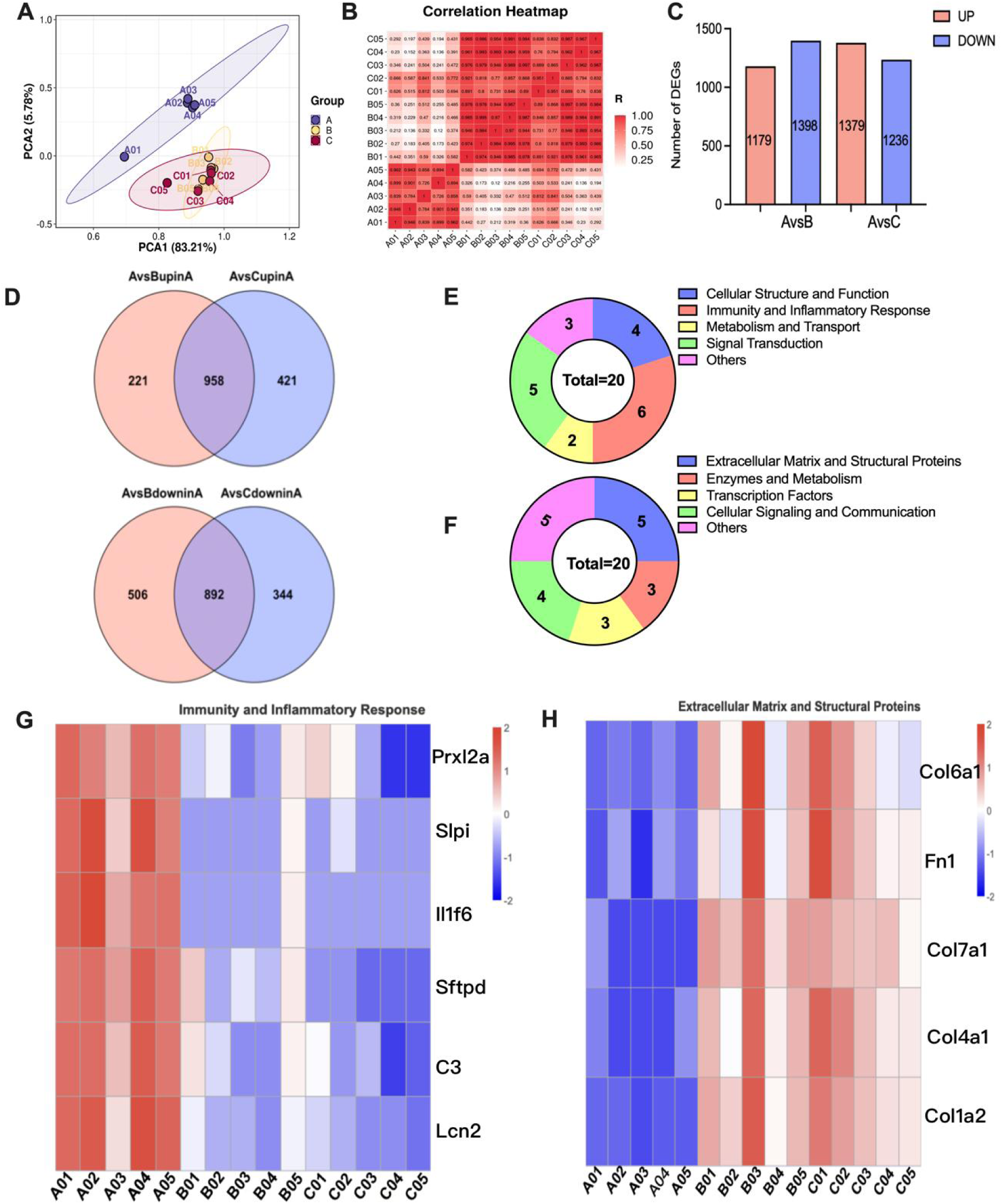
Transcriptome differences in uteri. (A) Principal component analysis (PCA) of uterus samples. (B) Correlation heat map between groups A to C. (C) Bar plots of differentially expressed genes (DEGs) in groups A to C. Red represents up-regulated DEGs; blue represents down-regulated DEGs. (D) Venn diagram of both up-regulated and down-regulated DEGs in group A compared to groups B and C. The overlapping regions represent genes that are co-up-regulated or co-down-regulated. (E) Donut chart of top 20 up-regulated and (F) down-regulated DEGs in group A. Different colors represent different gene function classifications in the Donut chart, and different numbers represent the number of DEGs in each functional category. (G) Heat map of immunity and inflammatory response, comparing groups A with B and C. (H) Heat map of extracellular matrix and structural proteins, comparing groups A with B and C. For the heat maps, red represents up-regulated DEGs; blue represents down-regulated DEGs. A, D1.5 recipients; B, D2.5 recipients; C, D3.5 recipients, pre-transfer blastocysts; E, post-transfer blastocysts. (n = 5).

Further analysis the expression of uterine DEGs, the top 20 genes that were significantly up-regulated in group A could be divided into 5 categories according to gene function: cellular structure and function (4/20), immunity and inflammatory response (6/20), metabolism and transport (2/20), signal transduction (5/20), and others (3/20) (Fig. 5E). The top 20 down-regulated genes in group A were mainly related to extracellular matrix and structural proteins (5/20), enzymes and metabolism (3/20), transcription factors (3/20), cellular signaling and communication (4/20), and others (5/20) (Fig. 5F). In the D1.5 uterus, some genes that promote inflammatory responses, such as *C3*, *Lcn2* and *Il1f6*, were significantly up-regulated (Fig. 5G). However, the genes related to the maintenance of cell structures in the collagen family were significantly down-regulated (Fig. 5H). Members of the collagen family played crucial roles in maintaining tissue integrity, providing structural support, and promoting intercellular interactions. The downregulation of their expression would result in a decline in the structural stability of uterine tissue.

### D1.5 uterine extracellular matrix remodeling was abnormal

ECM remodeling has an important impact on endometrium repair and regeneration, structure and function, and inflammatory response. GO and KEGG analyses were performed on the DEGs of A vs B and A vs C, respectively. The GO analysis revealed that the DEGs both in groups A vs B and A vs C were enriched in GO terms related to the composition and structure of the extracellular matrix (Fig. 6A, C). Meanwhile, the KEGG analysis indicated that the enriched pathways of A vs B and A vs C mainly were the HIF-1 signaling pathway, Focal adhesion, PI3K-Akt signaling pathway, Leukocyte transendothelial migration, and ECM-receptor interaction (Fig. 6B, D). Among them, the GSEA analysis showed that the ECM-receptor interaction pathway, which was enriched in the DEGs of A vs B and A vs C, was inhibited (Fig. 6E, F). Analysis DEGs of ECM-receptor interaction pathway in AvsB and AvsC revealed that there were 29 gene duplicates, which were mainly related to ECM structure maintenance (Fig. 6G, H). And the expression of collagen family genes related to the composition and structure of the extracellular matrix was significantly down-regulated (Fig. 6I). These results suggested that D1.5 uterine extracellular matrix remodeling was abnormal.

**Figure 6.**
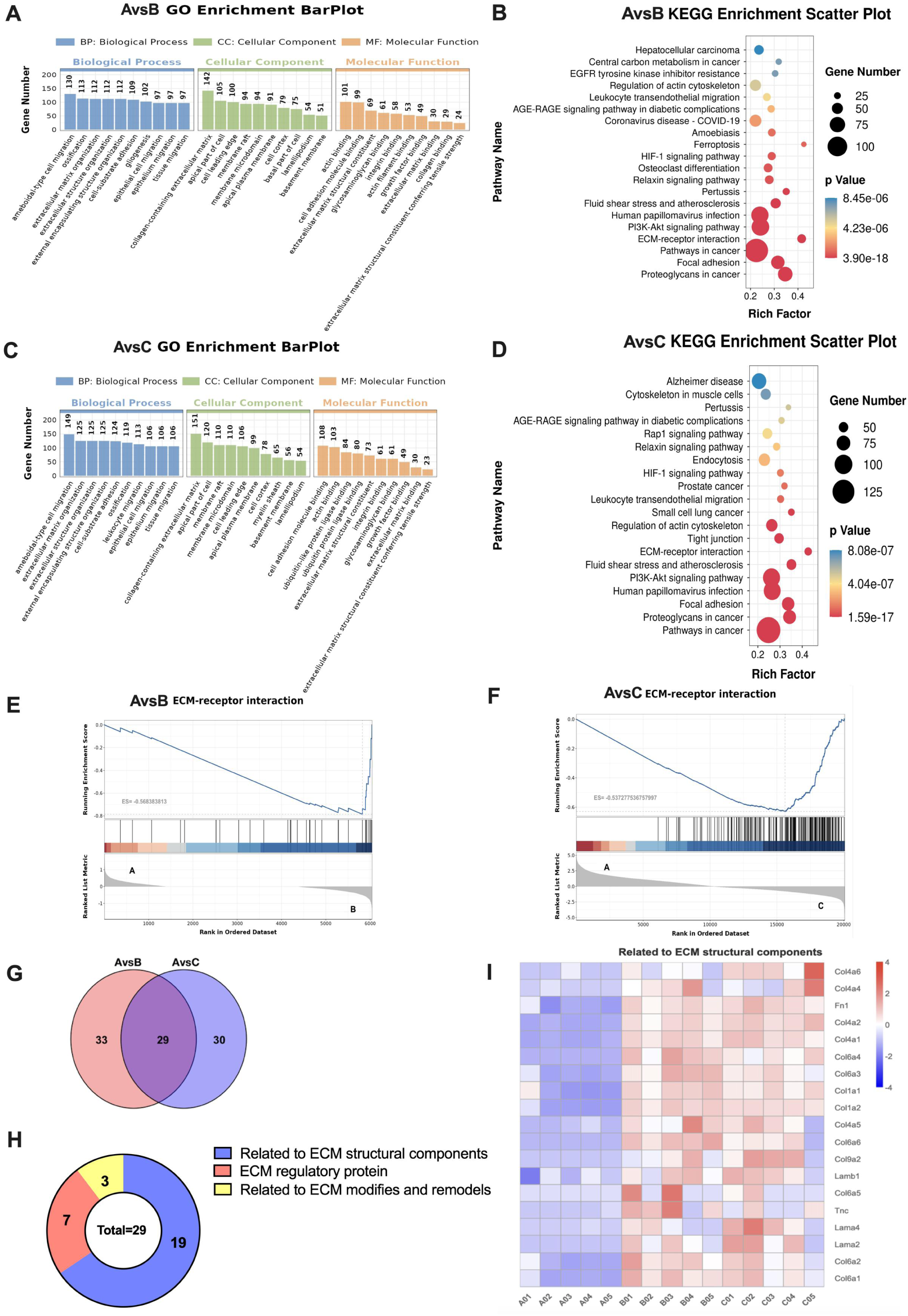
Extracellular matrix remodeling was abnormal in D1.5 uteri. (A, C) GO and KEGG analysis of DEGs between groups A and B. (B, D) GO and KEGG analysis of DEGs between groups A and C. For GO terms, each colored bar represents a different biological process, and the height of the bar indicates the number of DEGs involved. In the KEGG enrichment scatter plot, the color of the dot represents the p-value of the hypergeometric test; the smaller the value, the greater the reliability and statistical significance of the enrichment. The size of the dot represents the number of genes in the pathway; the larger the dot, the more DEGs are enriched in that pathway. (E, F) GSEA of ECM-receptor interaction between A vs B and A vs C. Negative ES values indicate that the pathway is inhibited in group A compared to the respective comparison group. (G) Venn diagram of DEGs in the ECM receptor interaction pathway between A vs B and A vs C. The overlapping regions represent genes that are co-downregulated in both comparisons. (H) Donut chart showing the distribution of ECM structural components among co-downregulated DEGs. (I) Heatmap of DEGs related to ECM structural components. Red represents up-regulated DEGs; blue represents down-regulated DEGs. A, D1.5 recipients; B, D2.5 recipients; C, D3.5 recipients. (n = 5).

### Immune and inflammatory dysregulation in the D1.5 uterus

Abnormal ECM remodeling may trigger or exacerbate inflammatory responses. To understand the gene expression trends, STEM analysis was conducted on all DEGs (with a cluster number of 8 and a *p*-value < 0.05). Four modules were significantly enriched (colored profiles 6, 1, 7, and 0), which included 198 DEGs (Fig. 7A). In profile 6, 78 DEGs showed a continuous increase in expression from D1.5 to D2.5 and remained unchanged at D3.5. In profile 1, 79 DEGs showed a continuous decrease in expression from D1.5 to D2.5 and remained unchanged at D3.5. In profile 7, 22 DEGs showed continuous upregulation. In profile 0, 19 DEGs showed a continuous down-regulation (Fig. 7B). GO and KEGG enrichment analyses were separately performed on the DEGs in profiles 6 and 1. KEGG analysis showed that in profile 6, the DEGs were mainly enriched in pathways such as ECM-receptor interaction, Focal adhesion, PI3K-Akt signaling pathway, Fluid shear stress and atherosclerosis, while GO analysis was mainly related to adhesion and the extracellular matrix (Fig. 7C, D). In profile 1, the DEGs were mainly enriched in pathways, such as the HIF-1 signaling pathway, Phagosome, ECM-receptor interaction, IL-17 signaling pathway, complement and coagulation cascades. GO terms were mainly enriched in immune response, inflammatory response, and the extracellular matrix (Fig. 7E, F). Both profile 6 and 1 were related to extracellular matrix, immune and inflammatory response. It was speculated that the D1.5 uterine environment was instability, accompanied by immune dysregulation.

**Figure 7.**
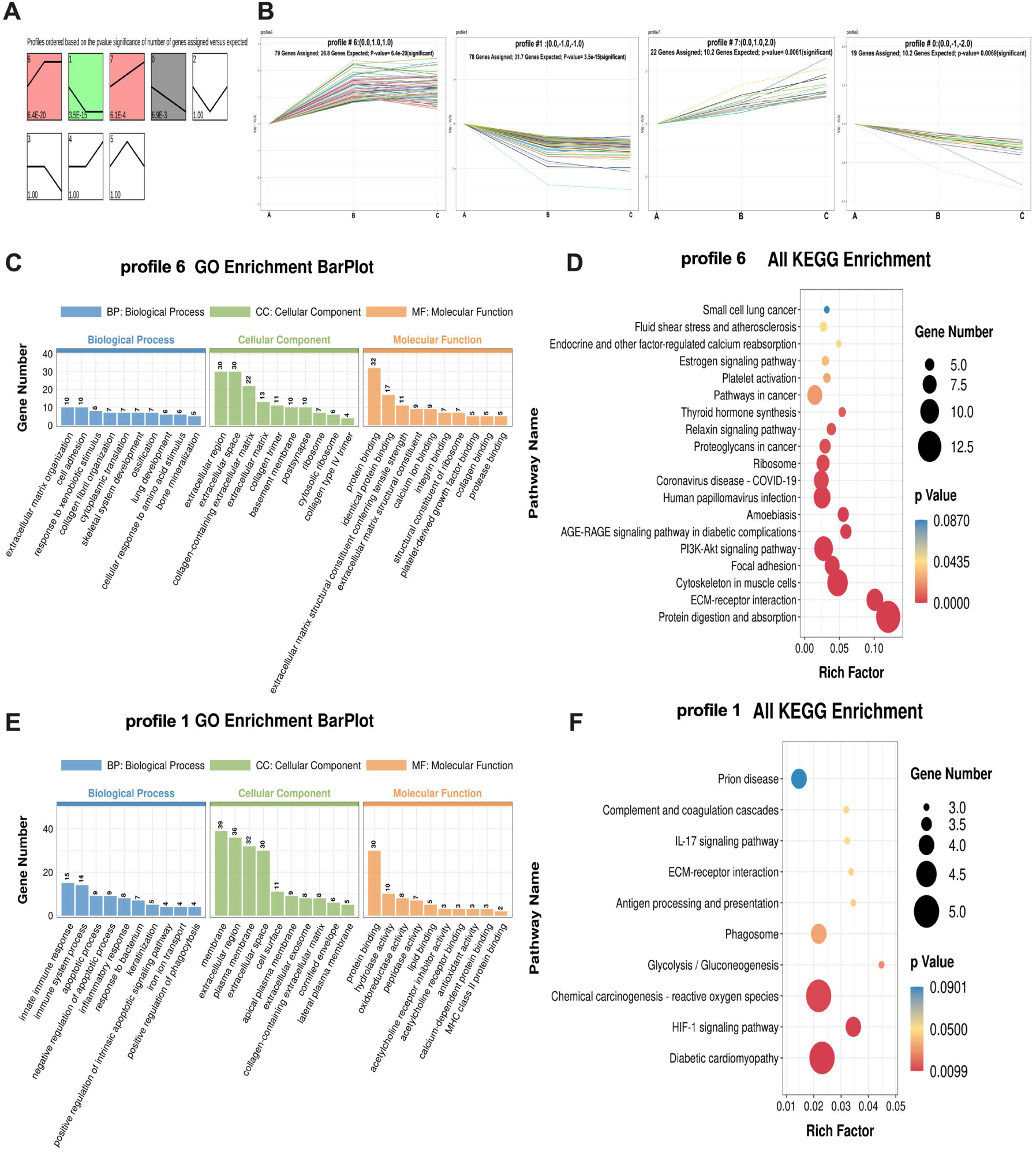
D1.5 uterine immune / inflammation dysregulation. (A) Trend analysis of DEGs in groups A, B and C. The colored modules represent significant trends. (B) Gene expression trends in profile 6, 1, 7, 0 modules. (C, D) GO and KEGG Analysis profile 6. (E, F) GO and KEGG analysis of profile 1. For GO term, each colored bar represents a different process and the height of the bar indicates the number of DEGs. And in KEGG enrichment scatter plot, the color of the dot represents the p value of the hypergeometric test, and the smaller the value, the greater the reliability and statistical significance of the test, the size of the dot represents the number of genes, and the larger the dot, the more DEGs in the pathway. A, D1.5 recipients; B, D2.5 recipients; C, D3.5 recipients. (n = 5).

### Screening of key genes regulating immunity and inflammation in the D1.5 uterus

Furthermore, clustering analysis on all genes should be conducted to clarify the state of the D1.5 uterus. After conducting WGCNA on genes with RPKM >1 and CV >= 0.0.5 (a minimum power = 5, kME > 0.7, and minimum module size = 30), we obtained a total of 13 modules. Among these, three modules (ME brown, ME yellow, and ME bule) were significantly correlated with group A (Fig. 8A). Further group correlation analysis and scatter plot of significant module-gene associations revealed that only the blue module showed significant differences (Fig. 8B, C). Enrichment analysis of the genes within the blue module indicated that they were primarily enriched in immune regulation-related pathways and GO terms such as Complement and coagulation cascades, Lysosome, Phagosome, and Leukocyte transendothelial migration, which is consistent with the results of the STEM analysis (Fig. 8D, E). Subsequently, Network interaction analysis was performed for genes with a degree greater than 40 in the blue module (Fig. 8F). The results showed that *Clca1*, *Smpdl3a*, *Tmprss4*, *Muc4*, *Cfb*, and *Nupr1* were crucial genes regulating immunity and inflammation in D1.5 uterus.

**Figure 8.**
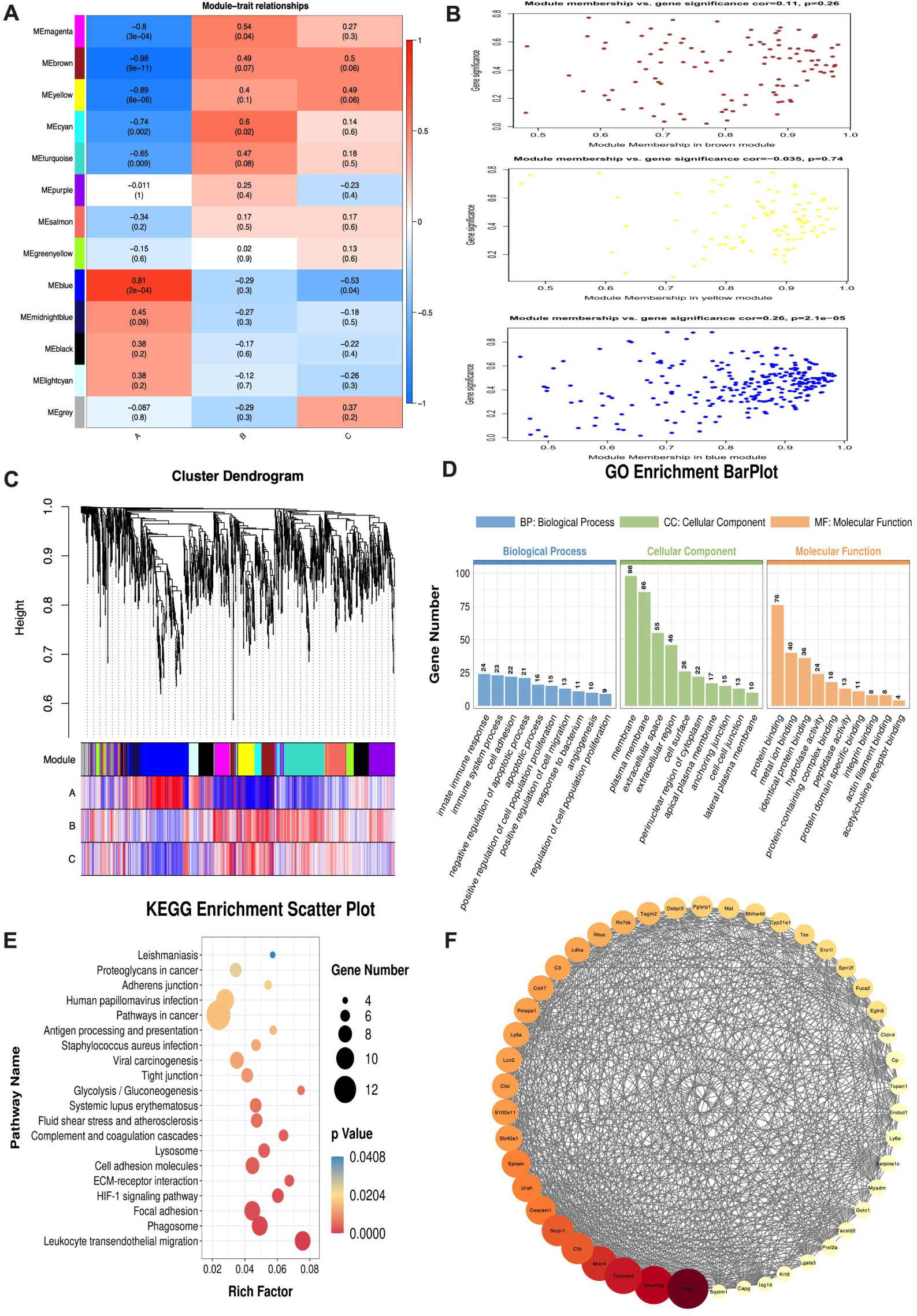
WGCNA analyse of D1.5 uterine. (A) Heatmap of module-trait ralationships. The horizontal axis represents the groups, while the vertical axis represents the modules. In the heatmap, the closer the color is to red, the stronger the positive correlation; conversely, the closer the color is to blue, the stronger the negative correlation. The values in the squares represent the correlation coefficients and p-values, respectively. (B) Scatter plot of significant module-gene associations in group A. Different colors represent different modules (brown、yellow and blue). The horizontal axis represents the kME values of the genes within the module, while the vertical axis represents the significance of the genes. Module members with higher cor values and smaller *p*-values are more representative of the module’s characteristics. (C) Hierarchical clustering dendrogram of genes and group correlation analysis. This figure is divided into three parts: The first part is the hierarchical clustering dendrogram of genes, which shows how genes are clustered based on their expression patterns.The second part displays the colors of the modules that the corresponding genes belong to, providing a visual representation of the gene-module associations.The third part shows the correlation between genes and their respective modules in each trait/group-related sample. The closer the color is to red, the stronger the positive correlation; conversely, the closer the color is to blue, the stronger the negative correlation. (D, E) GO and KEGG analysis of blue module in group A. For GO term, each colored bar represents a different process and the height of the bar indicates the number of DEGs. And in KEGG enrichment scatter plot, the color of the dot represents the *p*-value of the hypergeometric test, and the smaller the value, the greater the reliability and statistical significance of the test, the size of the dot represents the number of genes, and the larger the dot, the more DEGs in the pathway. (F) Gene Expression mmodule network diagram. The genes within the blue module are ranked based on their degree (refers to the sum of the connection strengths between a particular gene and other genes in the network). The genes with darker colors and larger dot sizes are considered hub genes. A, D1.5 recipients; B, D2.5 recipients; C, D3.5 recipients. (n = 5).

## DISCUSSION

Improving the success rate of embryo transfer remains a critical challenge that needs to be addressed in humans and animals. The success of embryo transfer depends on the quality of the embryos and the reproductive status of the recipients ^[23]^. High-quality embryos can significantly increase the success rate of a transfer, while low-quality embryos often result in lower pregnancy rates ^[24]^. Although mice have served as the classic model organism in biomedical research for over 60 years, embryo transfer technology has been studied extensively, but research on embryo transfer techniques remains inadequate. In this study, we first attempted to transfer well-developed blastocysts into the uteri of D1.5 recipient mice, but we failed to achieve successful implantation. Furthermore, when embryos were recovered from the uterus within 3 hours after transfer, it was found that the quality of the embryos had significantly declined, with a high proportion of abnormal embryos, reaching 68.67%. This discovery prompted us to reassess our previous understanding of embryo implantation failure. Traditionally, we tended to attribute the failure of embryo implantation solely to reduced uterine receptivity. However, the results of this study suggested that the underlying reasons may be far more complex than we initially thought.

The abnormal embryo staining results suggested that embryos with abnormal quality showed characteristics of dead cells. In order to clarify the specific dead reasons, we performed enrichment analysis on DEGs, which showed that the apoptotic pathway was activated. The apoptosis pathway is divided into two types: intrinsic and extrinsic. The intrinsic pathway is mainly caused by DNA damage and activation of P53^[25]^, while the extrinsic pathway is mainly caused by the death receptor pathway, including Fas, TNF-R1, TRAILR1and TRAIL-R2. After binding to their ligands, these death receptors activate intracellular signaling mediators, which ultimately trigger apoptosis ^[26]^. The results showed that the pro-apoptotic genes of both pathways were significantly up-regulated.

Further analysis of the differentially expressed genes is conducted to verify whether the intrinsic apoptotic pathway is induced by DNA damage. It is known that the uterus is a low-oxygen environment. When blastocysts are transferred into the uterus, they undergo a stress response^[27]^. Among the top 20 up-regulated genes in blastocysts, DNA damage-inducing transcription factor 4 (*Ddit4*) closely relates to DNA damage and regulates cell growth, proliferation, and survival by inhibiting the activity of the mammalian target of rapamycin protein complex 1 (mTORC1) ^[28]^. In tumor cell studies, the expression of the *Redd1* gene was significantly increased, and mTOR signaling was inhibited when hypoxia was induced ^[29]^. Although our study did not significantly enrich the mTOR signaling pathway, the significant up-regulation of *Ddit4* suggested that blastocyst cell function was somehow affected. Meanwhile, the down-regulation of Nsmce4a, which maintains the stability of chromatin structure and the mechanism of DNA damage repair, further confirmed that the blastocyst cell DNA had been damaged. Tubulin Beta 4B Class IVb (*Tubb4b*), a major component of microtubules, is essential for maintaining the cytoskeleton ^[30]^. Tubb4b-KO spermatogonia showed abnormal lysosomal membranes and cell morphology defects ^[31]^. All of these indicated that the embryos sustained damage to the cellular structure under the influence of the stress response. The originally orderly cell arrangement becomes disorganized, and the intercellular junctions become fragile. More seriously, this structural damage also led to DNA damage.

If DNA damage is not repaired promptly, it will affect normal life activities such as gene expression and protein synthesis, leading to serious disruptions in the transcription process. GO analysis showed that the DEGs were mainly enriched in cell signaling and cell cycle regulation. The coactivators *Taf3*, *Taf6l,* and *Taf10*, which are associated with the basic transcription factor pathway and play crucial roles in transcriptional regulation and cell growth and differentiation, were also down-regulated ^[32]^. Furthermore, GSEA analysis confirmed that the basic transcription factor pathway was inhibited. Cathepsin B (*Ctsb*) is believed to be involved in intracellular protein degradation and turnover ^[33]^, and it is upregulated in the apoptotic pathway and participates in the lysosomal pathway. It has been demonstrated that Ctsb promotes DNA damage and cell cycle arrest and induces lysosomal stress in retinoblastoma (RB) cells by inhibiting Brca1 expression and activating the STAT3/STING1 pathway ^[34]^. In our study, *Brca1* was also significantly downregulated (*p* = 0.044), which suggested that Ctsb may also promote DNA damage and cell cycle arrest by inhibiting the expression of *Brca1* in abnormal-quality blastocysts. What is even more noteworthy is that growth arrest and DNA damage-inducible 45 gamma (*Gadd45g*) is a stress-responsive protein involved in various biological processes, including DNA repair, cell cycle, and cell differentiation ^[35,36]^. It is also actively involved in the apoptotic pathway and has been shown to induce apoptosis in various cells, including tumor cells and cardiomyocytes. Therefore, when the DNA damage is not repaired, the blastocyst transcription is abnormal, and the intrinsic apoptotic pathway is activated.

It is undeniable that various cytokines and growth factors secreted by embryonic and uterine cells are necessary for the normal growth and development of the embryo. However, sperm entering the uterine cavity after normal mating may cause inflammatory responses ^[37]^. Studies have shown that *Slip* (secretory leukocyte protease inhibitor) is a serine protease inhibitor mainly secreted by mucosal epithelial cells in the respiratory, digestive, and reproductive tracts, and it exerts a regulatory effect on the inflammatory response ^[38–41]^. Its expression level is significantly up-regulated in the D1.5 uterus, so it can be inferred that the uterus at this time may experience inflammation due to semen residue. Additionally, due to the disordered expression levels of certain pro-inflammatory cytokines, such as *IL-1α* and *IL-1β*, in the uterus, embryos transferred into the uterus at this time are not only exposed to hypoxic stress but also to an altered uterine environment ^[42,43]^. Compared to the receptor uterus at D2.5 and D3.5, the pro-inflammatory factors *Il1f6* and *Lcn2* in the uterus at D1.5 are significantly up-regulated, actively participating in the regulation of inflammation ^[44,45]^. The KEGG analysis further suggested that the DEGs were significantly enriched in the TNF signaling pathway, a major mediator of apoptosis, inflammation, and immunity. It can be inferred that embryonic development will be affected in some manner due to the induction of various inflammatory factors. The blastocyst transcriptome analysis showed that TNF signaling pathways were also activated in abnormal-quality embryos, and the receptors *Tnfrsf12a* and *Tnfrsf11a* were also significantly up-regulated. Therefore, stress and DNA damage cause intrinsic apoptosis in embryos, and TNF also triggers apoptosis in embryos with abnormal quality through extrinsic pathways.

In addition to the impact of semen in the uterine cavity on D1.5, the environment of the uterus itself cannot be ignored. The theory of immune dysregulation, also known as the inflammation theory, was first proposed in the study of endometriosis due to local and systemic inflammation resulting from complex immune system dysregulation. When certain IL family chemokines and clusters of differentiation are highly expressed, inflammatory reactions occur ^[46,47]^. In the D1.5 uterus, *IL-1α*, *IL-1β*, *IL1F6*, *IL2*, *IL6*, and *IL17* were significantly up-regulated, and CDs such as CD40, CD44, and CD47 were also highly expressed. It has been found that high expression of inflammatory cytokines can trigger excessive inflammatory responses, which may adversely affect the success rate of in vitro fertilization-embryo transfer (IVF-ET) in infertile women ^[48]^. Moreover, the extracellular matrix (ECM), as a component of all organs, constantly undergoes remodeling to maintain tissue homeostasis and repair, playing a critical role in inflammatory reactions. Once immune cells penetrate the endothelium and enter the tissue, the ECM will guide the migration and regulate the survival and function of immune cells. It demonstrates that the ECM affects the homeostasis of immune cells in inflammatory tissues and that the immune environment influences the structure and remodeling of the ECM ^[49,50]^. Collagen, the most abundant structural protein in the ECM, provides the necessary support for tissues while guiding tissue development ^[51]^. However, COL1A2, COL4A1, COL7A1, COL6A1, and other related genes were significantly down-regulated in the D1.5 uterus. The GO analysis primarily highlighted terms related to the ECM, suggesting that the impaired ECM structure profoundly impacted uterine immune regulation. These two factors interacted, exacerbating the uterine environment.

Embryos are born to develop and grow in response to uterine secretions. Transcriptome analysis of the endometrium following asynchronous transfer in horses has shown that embryos can maintain pregnancy by extensively adapting their transcriptome in response to the asynchronous uterine state ^[52]^. When embryo quality is comparable, there is no significant difference between sequential embryo transfer (a two-step interval transfer of a cleavage-stage embryo followed by a blastocyst in one transfer cycle) and double-blastocyst transfer for women undergoing frozen-thawed embryo transfer cycles ^[53]^. However, in some clinical studies, the implantation rate of the day 6 blastocyst transfer group was significantly higher than the day 5 morula transfer group (27% vs. 12%, *p* < 0.001) ^[54]^. Similarly, pregnancy rates in asynchronous transfer studies in cattle and camels have been much lower than with synchronous transfer, and the lack of synchronization of the embryo with the reproductive tract in the early and middle stages may affect subsequent embryonic development ^[55–58]^. When 8-cell and blastocyst embryos are transferred to two uterine horns of the same recipient, the implantation window is regulated differently ^[59]^. Thus, although reproductive tract-embryo asynchrony affects pregnancy rates, embryos entering the uterus at different times have the potential to be regulated to adapt to the window of implantation. Furthermore, there may also be localized areas within the uterus that favor the development of some embryos to the detriment of others ^[55,60,61]^. In order to eliminate the effects of time asynchronization, we also attempted to transfer 2-cells into the D1.5 uterus, but the embryo was still unable to implant successfully (Supplemental Fig. 1). Our research confirms that no local area in the D1.5 uterus was favorable for embryonic development. This stage was not suitable for any period of embryo transfer. Moreover, the reason for embryo implantation failure is not due to a lack of synchronization between endometrial development and embryonic development; rather, it is primarily caused by uterine immune dysregulation that adversely affects the uterine environment, thereby hindering embryonic development. We plan further to elucidate the specific regulatory mechanisms in future research.

In summary, immune dysregulation and residual semen within the uterine environment during embryo transfer to the D1.5 recipient uterus results in blastocyst stress and inflammation. On the one hand, they disrupt normal DNA damage repair, activating the intrinsic apoptosis pathway. On the other hand, they facilitate the activation of the extrinsic apoptotic pathway via death receptor TNFR, simultaneously promoting the death of blastocyst cells. Consequently, the embryo could not further develop towards the implantation window, ultimately causing implantation failure. However, further in-depth exploration and research are still required to elucidate the mechanisms underlying this immune dysregulation and its precise impact on embryonic survival. Furthermore, this study warns us that immune dysregulation in the uterine environment is also a critical factor affecting pregnancy rates in assisted reproduction(Supplemental Fig. 2).

## Acknowledgements

The authors are grateful to Geekgene (Geekgene Technology Co., Ltd., Beijing, China) for their MultiBarcode single-cell transcriptome sequencing services.

