## Supplemental Fig. 1 &Supplemental Fig. 2 for "Uterine immune dysregulation after asynchronous transfer induced embryonic apoptosis in mice"

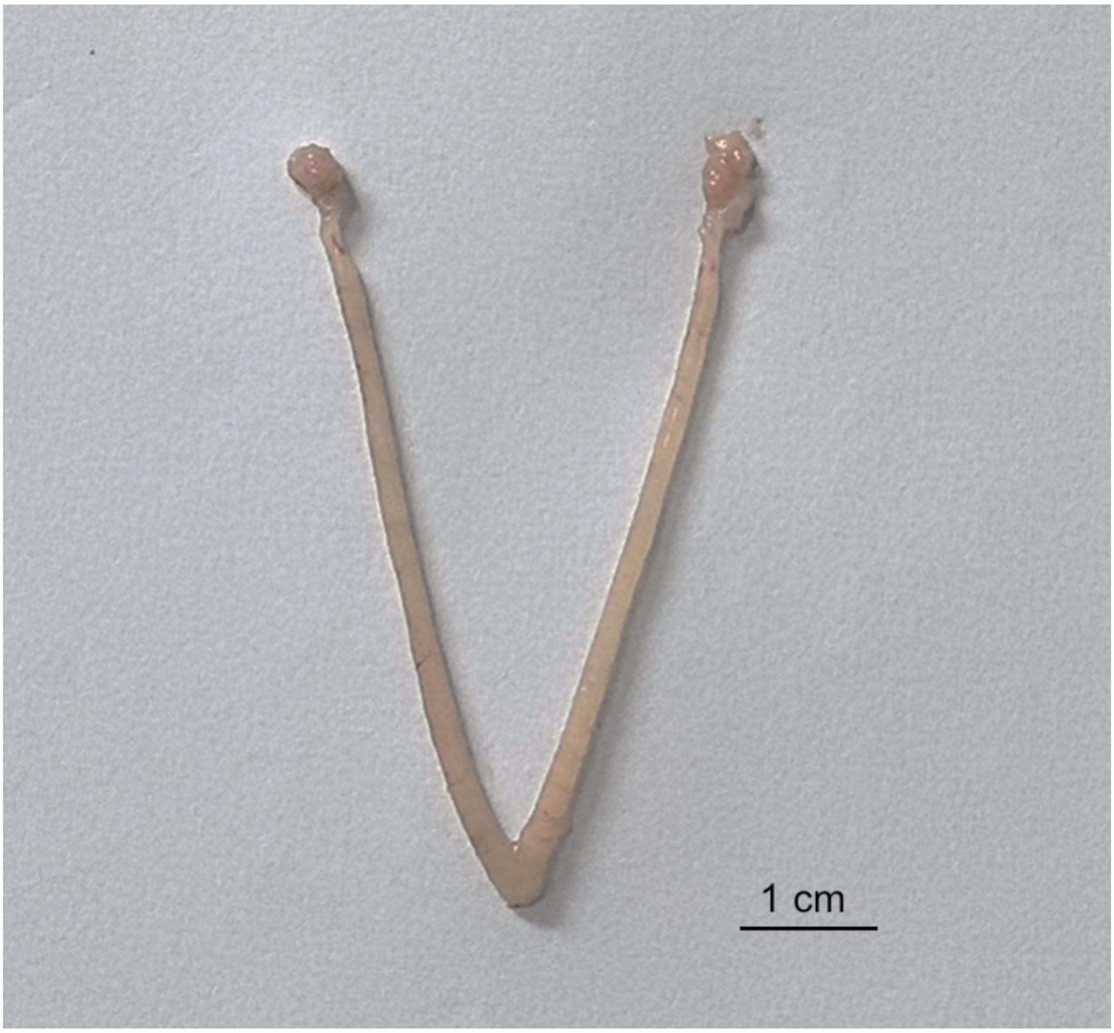


**Supplemental figure1. Implantation sites of embryos on Day7.5**. Eight 2-cell were transferred into D1.5 recipient's uterus. (bar = 1 cm).

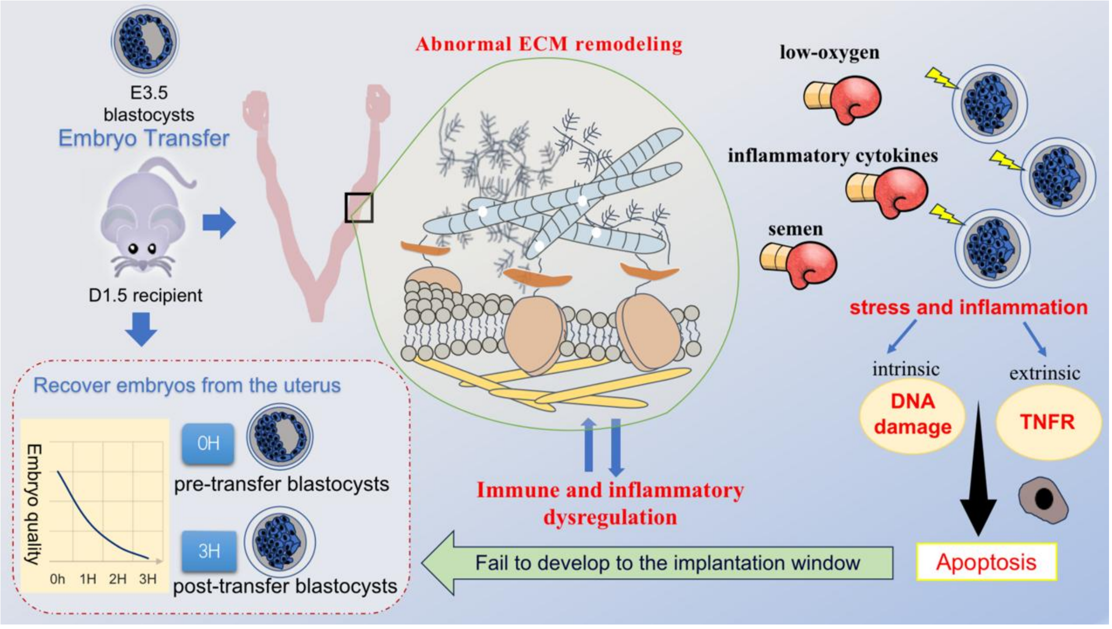


**Supplemental figure2. Graphical Abstract of the study.** When blastocysts are transferred into D1.5 recipient mice, the embryos experience stress and inflammation due to dysregulation of the immune system in the uterus. On the one hand, they disrupt normal DNA damage repair, activating the intrinsic apoptosis pathway. On the other hand, they facilitate the activation of the extrinsic apoptotic pathway via the death receptor TNFR, simultaneously promoting the death of blastocyst cells. Consequently, the embryo could not further develop towards the implantation window, ultimately causing implantation failure.
